# A virion-based assay for glycoprotein thermostability reveals key determinants of filovirus entry and its inhibition

**DOI:** 10.1101/2020.02.25.965772

**Authors:** Robert H. Bortz, Anthony C. Wong, Michael G. Grodus, Hannah S. Recht, Marc C. Pulanco, Gorka Lasso, Simon J. Anthony, Eva Mittler, Rohit K. Jangra, Kartik Chandran

## Abstract

Ebola virus (EBOV) entry into cells is mediated by its spike glycoprotein (GP). Following attachment and internalization, virions traffic to late endosomes where GP is cleaved by host cysteine proteases. Cleaved GP then binds its cellular receptor, Niemann-Pick C1. In response to an unknown cellular trigger, GP undergoes conformational rearrangements that drive fusion of viral and endosomal membranes . The temperature-dependent stability (thermostability) of the pre-fusion conformers of ‘Class I’ viral fusion glycoproteins, including those of filovirus GPs, has provided insights into their propensity to undergo fusion-related rearrangements. However, previously described assays have relied on soluble glycoprotein ectodomains. Here, we developed a simple ELISA-based assay that uses the temperature-dependent loss of conformational epitopes to measure thermostability of GP embedded in viral membranes. The base and glycan cap subdomains of all filovirus GPs tested suffered a concerted loss of pre-fusion conformation at elevated temperatures, but did so at different temperature ranges, indicating virus-specific differences in thermostability. Despite these differences, all of these GPs displayed reduced thermostability upon cleavage to GP_CL_. Surprisingly, acid pH enhanced, rather than decreased, GP thermostability, suggesting it could enhance viral survival in hostile endo/lysosomal compartments. Finally, we confirmed and extended previous findings that some small-molecule inhibitors of filovirus entry destabilize EBOV GP and uncovered evidence that the most potent inhibitors act through multiple mechanisms. We establish the epitope-loss ELISA as a useful tool for studies of filovirus entry, engineering of GP variants with enhanced stability for use in vaccine development, and discovery of new stability-modulating antivirals.

**Importance:** Though a vaccine for Ebola virus has been approved by the FDA within the past year, no FDA-approved therapeutics are available to treat infections by Ebola virus or other filoviruses. The development of such countermeasures is challenged by our limited understanding of the mechanism by which Ebola virus enters cells, especially at the final step of membrane fusion. The sole surface-exposed viral protein, GP, mediates key steps in virus entry, including membrane fusion, and undergoes major structural rearrangements during this process. The stability of GP at elevated temperatures (thermostability) can provide insights into its capacity to undergo these structural rearrangements. Here, we describe a new assay that uses GP-specific antibodies to measure GP thermostability under a variety of conditions relevant to viral entry. We show that proteolytic cleavage and acid pH have significant effects on GP thermostability that shed light on their respective roles in viral entry. We also show that the assay can be used to study how small-molecule entry inhibitors affect GP stability. This work provides a simple and readily accessible assay to engineer forms of GP with enhanced stability that could be useful as part of an antiviral vaccine and to discover and improve drugs that act by modulating the stability of GP.

## Introduction

Ebola virus (EBOV) is an enveloped negative-strand RNA virus in the family *Filoviridae*. The virus is responsible for causing Ebola virus disease (EVD), a devastating clinical syndrome that is characterized by early nonspecific findings followed by severe gastrointestinal symptoms and hemorrhage complications (1–3). Multiple ebolaviruses, including Sudan virus (SUDV), are capable of causing human disease with significant mortality. However, EBOV has been responsible for the majority of recorded human outbreaks including two recent large-scale outbreaks—the unprecedented 2013-2016 West African epidemic (4), and an ongoing outbreak in the Democratic Republic of the Congo (5). Although an EBOV vaccine was recently approved by the FDA (6), no FDA-approved therapeutics are currently available for any of the filoviruses (7).

EBOV entry into cells requires a complex sequence of events that are mediated by the sole surface-exposed viral glycoprotein (GP). GP is composed of two subunits that are tethered by non-covalent interactions and an intersubunit disulfide bond. GP1, the membrane-distal subunit, contains the receptor-binding site (RBS) which is shielded by the glycan cap, and a variable and highly glycosylated mucin-like domain (Muc) (8–12). GP2, the transmembrane subunit, mediates membrane fusion and contains sequences characteristic of Class I membrane fusion proteins, including an internal fusion loop, N– and C–terminal heptad repeats that form α–helical coiled coils, and a flexible membrane-proximal extracellular region (8, 13, 14). Virions initially attach to host cells through both GP and viral membrane-mediated interactions with multiple cellular attachment factors (15, 16). After internalization via a macropinocytosis-like mechanism (17–19), virions traffic along the endocytic pathway to late endo/lysosomal compartments (20, 21). Here, they are cleaved by host proteases cathepsins B and L (CatB and CatL, respectively) in the β13–14 loop of GP1. This removes the GP1 glycan cap and Muc, exposing the RBS and producing a metastable, primed intermediate of GP, GP_CL_ (22–24). GP_CL_ is then able to bind to the host receptor, Niemann-Pick C1 (NPC1), a cholesterol transporter located in the endo-lysosomal membrane. Binding to NPC1 is necessary but not sufficient to facilitate viral entry (25, 26).

Despite the considerable body of research on EBOV entry, its precise mechanism, especially at the membrane fusion step, remains poorly understood. Following receptor binding, an unknown trigger causes a series of conformational rearrangements that bring the host endosomal and viral membranes together to form a fusion pore, enabling the release of the viral nucleocapsid into the cytoplasm. Although the intermediate conformations of GP during membrane fusion have not been experimentally visualized, GP2 rearrangements culminate in a post-fusion 6-helix bundle similar to that observed for other Class 1 viral fusogens, suggesting an analogous fusion mechanism (13, 14).

The thermostability of Class I fusion glycoproteins has provided a valuable surrogate for their capacity to undergo entry-related conformational changes. For example, mutations in the influenza A virus and human immunodeficiency virus glycoproteins that alter their thermostability and viral infectivity also affect their propensity to undergo acid pH- or receptor-mediated conformational changes during membrane fusion (27–29). Similarly, the stability of EBOV GP can impact viral infectivity. Specifically, a GP1 mutation, A82V, that emerged during the West African outbreak decreased GP thermostability and increased viral infectivity in cell culture (30, 31). By contrast, small molecule inhibitors bind into a pocket at the base of GP and destabilize it (32–34), indicating that reductions in GP thermostability can have opposing effects on EBOV entry and infection. Further, we recently showed that a thermostabilizing GP mutation, R64A, abolishes infection and that compensatory second-site mutations reduce GP thermostability (35). Work to date thus indicates that both decreased and increased GP thermostability can influence EBOV entry.

Although informative, previous studies of EBOV GP thermostability have largely relied on either hydrophobic dye- (32–34) or membrane-binding assays (36) with recombinant glycoprotein ectodomains. In the former type of assay, region-specific conformational changes in GP cannot be readily assessed, whereas in the latter, the potential stabilizing effects of the GP transmembrane domain and the membrane environment are neglected. Here, we sought to bridge the gap between these approaches by developing a quantitative assay for heat-induced conformational changes in full-length GP displayed on the membranes of intact vesicular stomatitis virus (VSV) particles. Specifically, we used a panel of GP-specific conformation-sensitive monoclonal antibodies (mAbs) to detect heat-induced loss of the pre-fusion conformation of EBOV GP under different conditions and extended these observations to other filovirus glycoproteins. We show that the structural core of filovirus GP undergoes a concerted conformational rearrangement at a characteristic temperature range that is lowered by GP proteolytic cleavage and by some, but not all, mutations that modulate its proteolytic susceptibility. Counterintuitively, we find that GP thermostability is increased by acid pH. Finally, we confirm that some selective estrogen receptor modulators (SERMs) can destabilize EBOV GP, as shown previously with recombinant GP ectodomains, but find that their destabilizing effect on filovirus glycoproteins can be at least partially decoupled from their antiviral activity.

## Results

### Development of an epitope-loss ELISA to study the stability of the EBOV GP

To interrogate the thermostability of native, full-length EBOV GP in biological membranes, we developed an assay that utilizes conformation-specific mAbs to detect the heat-induced loss of structure of a given epitope. Recombinant VSVs expressing EBOV GP (rVSV-GP) were heated at a range of temperatures, cooled to 4°C, and directly coated onto ELISA plates. The binding capacities of selected conformation-specific mAbs were then determined by ELISA. We also tested the thermostability of both a fully infectious mutant lacking the mucin-like domain (Muc), GP_Δmuc_ (22, 37). We first used the well-characterized EBOV GP-specific mAb KZ52, which detects a conformational GP1–GP2 intersubunit epitope in the pre-fusion GP trimer (that is maintained in GP_ΔMuc_) (8, 38). The thermal denaturation curves obtained for GP and GP_ΔMuc_ showed a similar sigmoid shape: strong KZ52 binding was observed at low temperatures but decreased to background levels between 56–64°C (Fig 1A). This profile is consistent with a two-state model in which the KZ52 epitope is either present or absent in GP, with an increasing probability of denaturing the epitope with increasing temperature.

**Fig 1:**
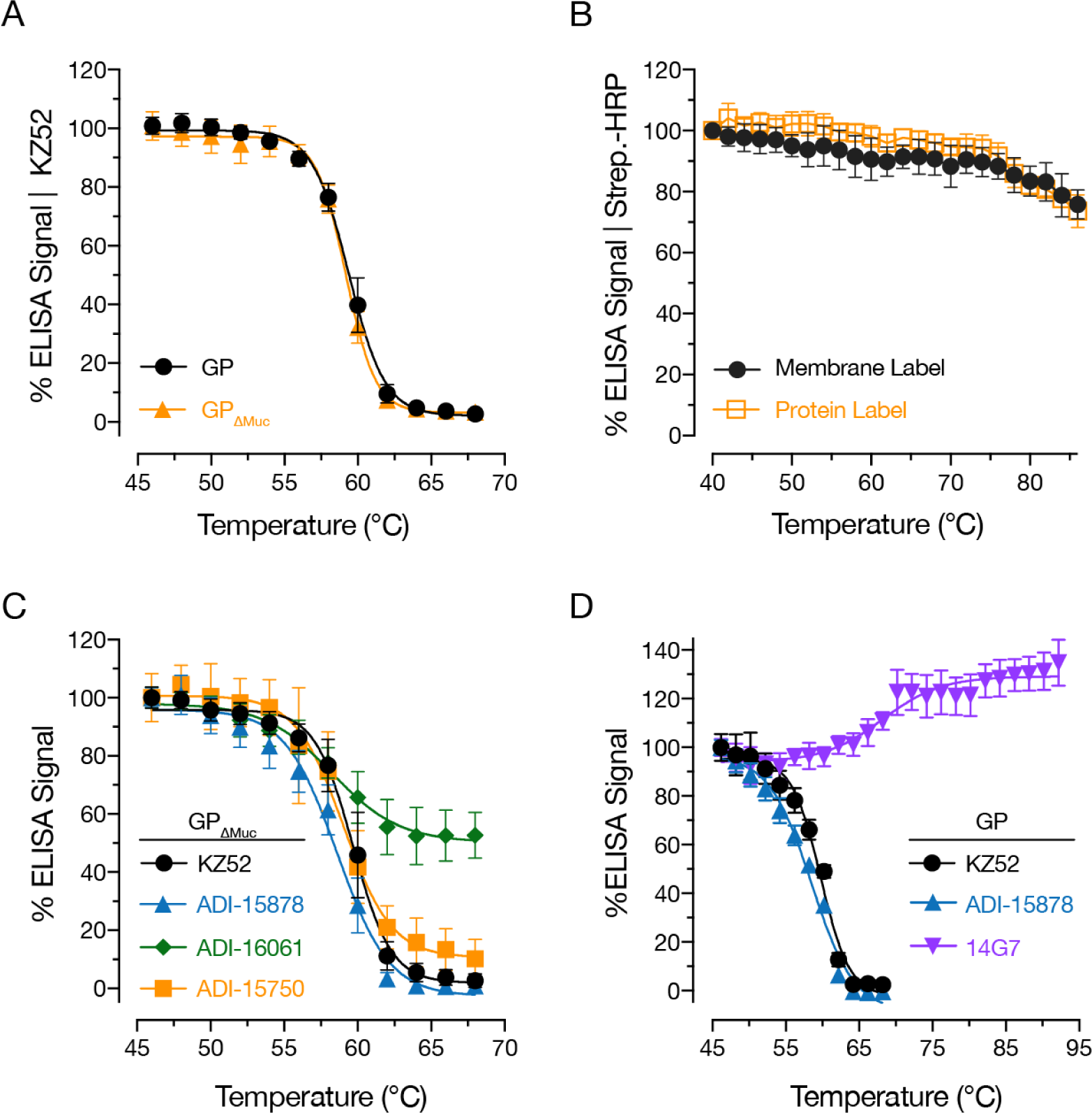
Thermal denaturation curves for pre-fusion epitopes in uncleaved EBOV GP. (A) rVSV-GP and rVSV-GP_ΔMuc_ were incubated at the indicated temperatures for 10 min, after which the samples were cooled to 4°C and KZ52 binding was assessed by ELISA. Averages±SD, n=9 from 3 independent experiments. (B) Membrane- and protein-labeled rVSV-GP_ΔMuc_ preparations were incubated at the indicated temperatures and biotin-labeled particles were detected with streptavidin-HRP by ELISA. Averages±SD, n=6 from 2 independent experiments. (C) Effect of virus pre-incubation at the indicated temperatures on binding by mAbs ADI-15878, ADI-16061, ADI-15750, and KZ52. Averages±SD, n=15 from 4 independent experiments (except ADI-15878, n=12 from 4 independent experiments). (D) Effect of virus pre-incubation at the indicated temperatures on binding by Muc-specific mAb 14G7. Binding curves for KZ52 and ADI-15878 binding are shown for comparison. Averages±SD, n=9 from three independent experiments.

Similar half-maximal mAb binding temperatures, (T_m_ ≍ 59℃) were obtained for GP and GP_Δmuc_, suggesting that Muc does not contribute to the stability of the GP pre-fusion conformation. Therefore, we largely used viral particles bearing GP_ΔMuc_ in the following experiments.

To rule out the trivial possibility that the loss of KZ52 binding was due to decreased virion capture onto ELISA plates or shedding of GP from viral particles at elevated temperatures, protein- or membrane-biotinylated preparations of VSV-GP_ΔMuc_ were also subjected to the same protocol, and the virion-associated biotin signal was measured by ELISA. Both GP and viral particles were detected at all temperatures tested, including temperatures far exceeding those used in the epitope loss ELISA (Fig 1B). Importantly, only minimal reductions in biotin signal were observed at temperatures over which KZ52 binding titrated (56–64°C), with significant decreases only occurring at temperatures >72°C (Fig 1B). These experiments indicate that the irreversible loss of KZ52 binding to GP at elevated temperatures is a consequence of the thermal denaturation of the KZ52 epitope and not the loss of viral particles or GP.

### The GP base and glycan cap subdomains undergo a concerted loss of pre-fusion conformation at elevated temperatures

To investigate if the loss of the KZ52 epitope at elevated temperatures was specific to this epitope or instead reflected larger-scale changes in the pre-fusion conformation of GP, we tested additional conformation-sensitive mAbs whose epitopes are distinct from that of KZ52: ADI-15750, ADI-15878 and ADI-16061 (39). ADI-15750 recognizes the glycan cap subdomain in GP; ADI-15878 binds a distinct interprotomer epitope in the GP base spanning GP1 and the GP2 fusion loop; and ADI-16061 recognizes a GP2 epitope in the stalk of the GP trimer (39). The thermal denaturation curves for the ADI-15750 and ADI-15878 epitopes were superimposable with that of KZ52, with T_m_ values of ∼59℃(Fig 1C). By contrast, the ADI-16061 epitope was largely resistant to elevated temperatures, possibly because this epitope in the GP2 HR2 domain is stabilized by its proximity to the GP membrane anchor. Alternatively, it is possible that the ADI-16061 epitope partially renatures during the cooling step or subsequent steps in the ELISA (Fig 1C).

In contrast to the GP base and glycan cap subdomains probed above, Muc was shown to be largely disordered, with several Muc-specific mAbs recognizing linear epitopes (11, 12, 40–42). To investigate the thermostability of Muc, we used mAb 14G7, which recognizes a linear Muc epitope (40). Over the temperature range at which the base and glycan cap epitopes were lost, we observed no appreciable reduction in the 14G7 epitope (Fig 1D). Instead, 14G7 recognition was enhanced at very high temperatures (Fig 1D), possibly due to the increased exposure of its linear epitope (40). Taken together, these experiments demonstrate that the highly structured regions of the GP trimer, including the base and glycan cap subdomains, suffer a concerted, irreversible loss of their pre-fusion conformation at elevated temperatures.

### EBOV GP is destabilized by proteolytic cleavage

Proteolytic cleavage of GP by endosomal cysteine cathepsins CatB and CatL exposes the binding site for its critical endo/lysosomal receptor, NPC1, and primes it to undergo subsequent entry-related conformational changes (36, 43, 44). Previous work also suggests that cleaved GP conformers (GP_CL_) generated *in vitro* are more conformationally labile than their uncleaved counterparts (36, 44). To investigate the consequences of proteolytic processing on GP thermostability, rVSV-GP was incubated with thermolysin (THL) as described previously and tested in the epitope-loss ELISA (23, 44). THL is proposed to mimic the cleavage of GP by CatB during viral entry (23). GP cleavage was verified by western blot (Fig 2A). Although the thermal denaturation curve for THL-cleaved GP_CL_ (GP_THL_) resembled those of GP and GP_ΔMuc_ in sigmoidal shape, it was left-shifted by ∼6℃relative to the latter, indicating decreased stability (Fig 2B and C). We obtained similar findings with the GP base epitope of mAb ADI-15878 and the RBS epitope of MR72 (45, 9). ADI-16061’s GP stalk epitope was even more resistant to denaturation in GP_THL_ than in its uncleaved counterpart (Fig 2D).

**Fig 2:**
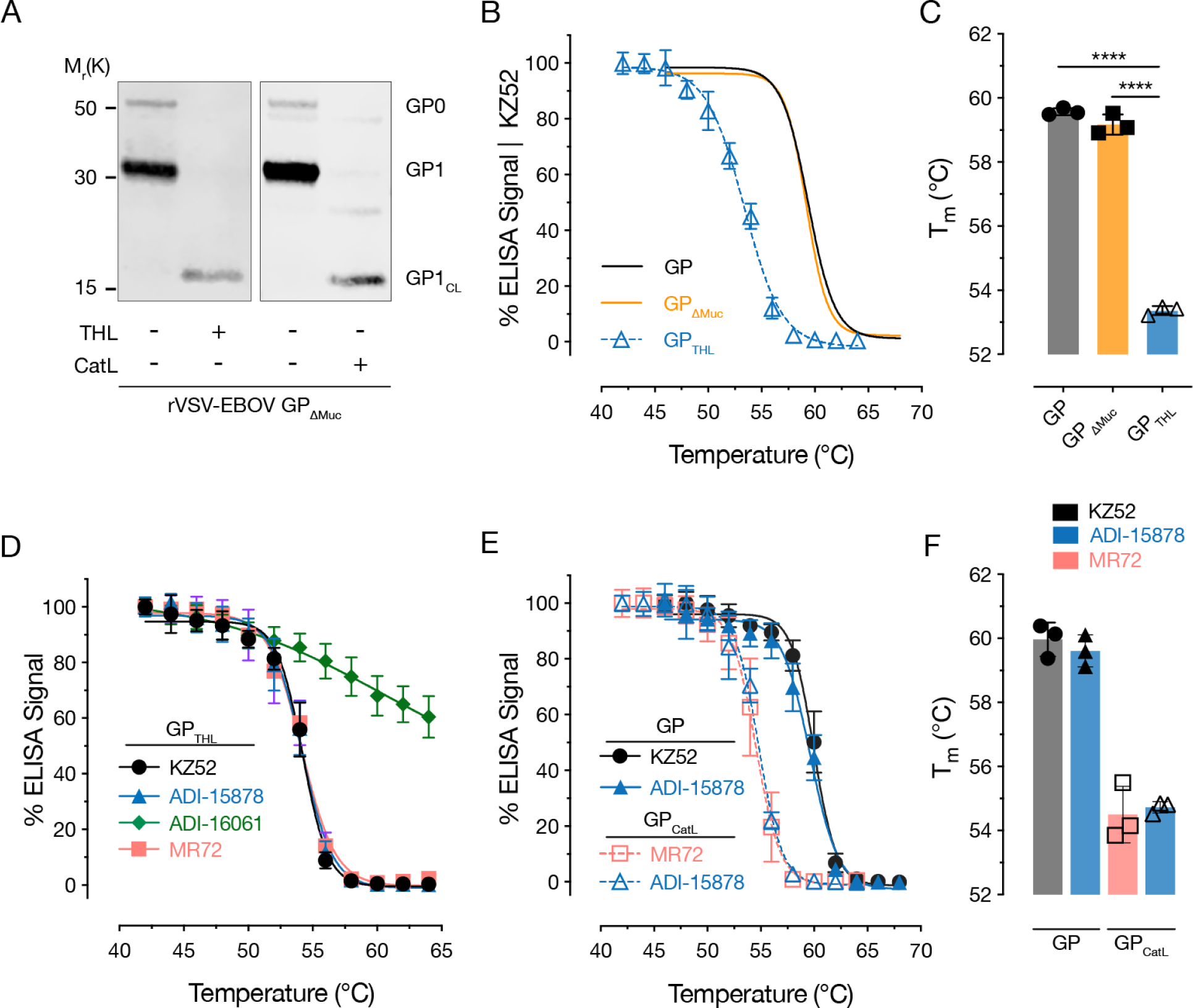
Proteolytic cleavage with thermolysin (THL) and cathepsin L (CatL) destabilizes EBOV GP. (A) THL and CatL cleavage of rVSV-GP_ΔMuc_ was verified by western blot with anti-GP1 N terminal peptide polyclonal rabbit sera. (B–C) Thermal denaturation curves (B) and calculated melting temperature values (T_m_) (C) obtained with KZ52 for rVSVs bearing GP_THL_ compared to uncleaved GP and GP_ΔMuc_. Data for GP and GP_ΔMuc_ are from Fig 1 and are shown for comparison. (D) Thermal denaturation curves for rVSV-GP_THL_ obtained with mAbs ADI-15878, ADI-16061 and MR72. Averages±SD, n=15 from 5 independent experiments except for MR72 and ADI-15878 (n=12 from 4 independent experiments). (E–F) Thermal denaturation curves (E) and T_m_ values (F) for GP and GP_CatL_ as assessed with KZ52 and ADI-15878 (GP) and MR72 and ADI-15878 (GP_CatL_). Averages±SD, n=9 from three independent experiments.

We next asked if cleavage with the more entry-relevant endosomal cysteine protease CatL also destabilizes GP. Accordingly, we treated rVSV-GP with recombinant human CatL and verified the generation of CatL-cleaved GP_CL_ (GP_CatL_) by western blot (Fig 2A). Because KZ52 does not bind GP_CatL_ (46), we used ADI-15878 and MR72 to assess its thermostability (46). As expected, CatL cleavage also destabilized GP and did so to a similar extent as THL, with a T_m_ of ∼55℃for both ADI-15878 and MR72 (Fig 2E and F). Together, these experiments indicate that the proteolytic removal of the glycan cap and cleavage in the β13–14 loop of GP1 associated with GP◊GP_CL_ cleavage destabilize the pre-fusion conformation of GP, affording one mechanism by which GP is primed for membrane fusion-related conformational changes during entry.

### The GP proteins of other ebolaviruses are also destabilized by proteolytic cleavage

Because GP→GP_CL_ cleavage is a prerequisite for cell entry by all filoviruses that have been evaluated to date (22, 24, 47), we postulated that this cleavage also destabilizes GPs from divergent ebolaviruses. Accordingly, we used the pan-ebolavirus base-binding mAb ADI-15878 to probe the thermal stability of GPs from the divergent ebolaviruses Sudan virus (SUDV) and the recently discovered Bombali virus (BOMV) in the epitope-loss ELISA (48–51). THL cleavage of SUDV GP and BOMV GP was verified by western blot (Fig 3A). As described for other ebolaviruses, BOMV GP1 was cleaved by THL to a ∼17K product (GP_17K_), which is consistent with its capacity to recognize NPC1 through exposure of its RBS (51). Like EBOV GP_THL_, SUDV and BOMV GP_THL_ displayed reduced themostability relative to their uncleaved counterparts, indicating that GP cleavage plays a similar role for several (and likely, all) ebolaviruses in destabilizing the pre-fusion conformation of GP (Fig 3B and C). Although SUDV GP resembled EBOV GP in thermostability in both its uncleaved and cleaved forms, BOMV GP was much less stable. Indeed, uncleaved BOMV GP had a similar T_m_ to that of EBOV GP_CL_ (Fig 3B and C). These findings reveal differences in the intrinsic stability of the GP pre-fusion conformation among ebolaviruses, with potential implications for their cell entry and infection mechanisms.

**Fig 3:**
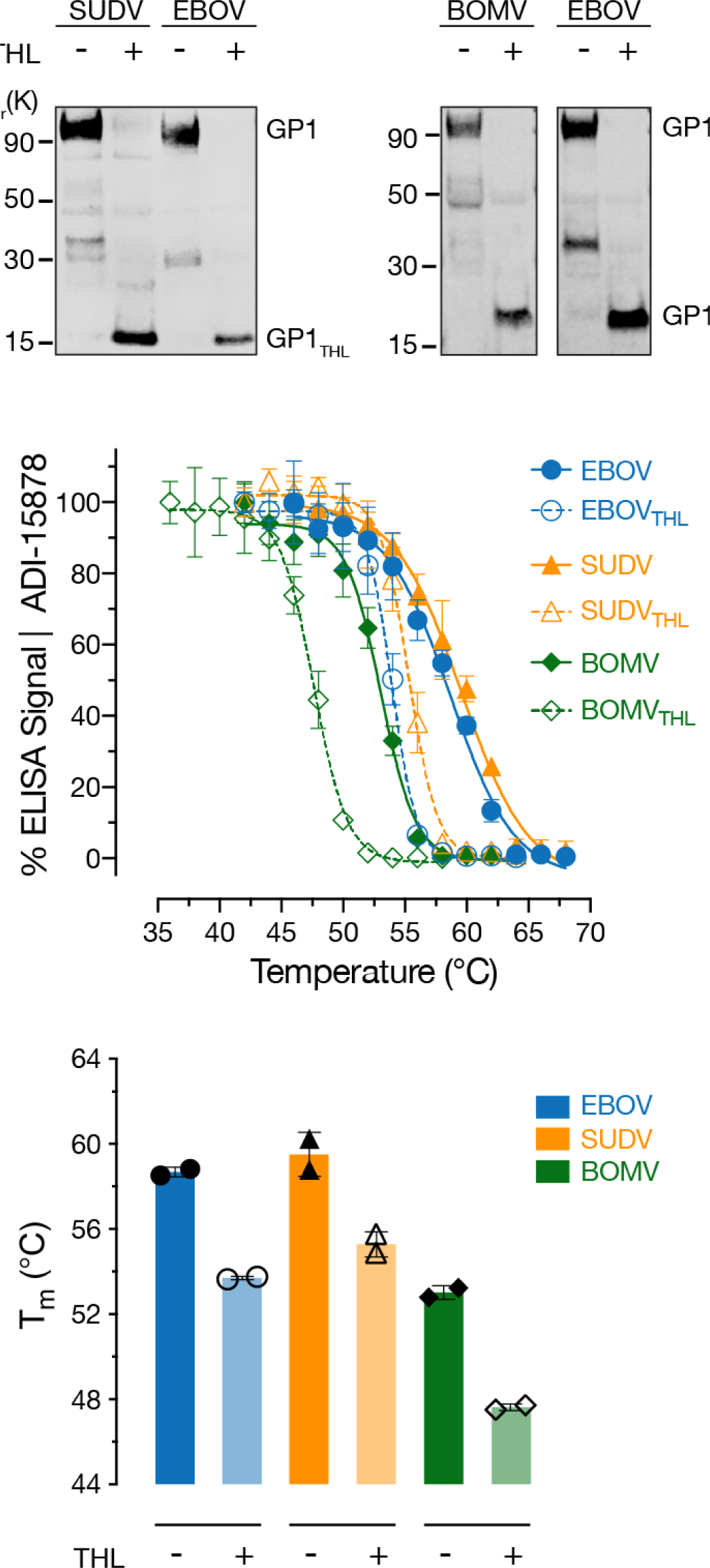
Thermostability of GPs from other ebolaviruses. (A) THL cleavage conditions for rVSV-SUDV GP and rVSV-BOMV GP were verified by western blot with anti-GP1 N terminal peptide polyclonal rabbit sera. (B) Thermal denaturation curves for rVSVs bearing GP and GP_THL_ from EBOV, SUDV and BOMV were determined with ADI-15878. Averages±SD, n=6 from 2 independent experiments (C) Calculated T_m_ values from the thermal denaturation curves in panel B. Averages±SD, n=2 from 2 independent experiments.

### A subset of CatB-independent EBOV GP mutants exhibits decreased thermostability

In a previous study, we selected and characterized EBOV GP mutants bearing single point mutations that afforded CatB-independent virus entry (44). Because two of these mutations (I584F and K588R) were located at the GP1-GP2 intersubunit interface, we proposed that they destabilize the GP pre-fusion conformation in a manner that enables virions to bypass the CatB cleavage requirement. To more rigorously evaluate this hypothesis, we assessed the thermostability of the GP_ΔMuc_ and GP_THL_ forms of the CatB-independent N40K, D47V, I584F and K588R mutants in the epitope-loss ELISA (Fig 4). We observed little or no left-shift in the thermal denaturation curves for GP(N40K) and GP(D47V), indicating that these mutations do not confer CatB independence by destabilizing GP (Fig 4A and D). By contrast, the thermal denaturation curves of both GP(I584F) and GP(K588R) and their GP_THL_ intermediates were left-shifted relative to those of GP(WT). Unexpectedly, K588R rendered GP considerably more unstable than did I584F, despite the fact that the latter was much more sensitive than the former to proteolytic degradation (Fig 4B, C and D) (44). These results lend further support to the hypothesis that the key function of GP cleavage by CatB is to destabilize GP’s pre-fusion conformation (35, 36, 44). They also indicate, however, that some GP mutants bypass the CatB cleavage requirement by mechanisms other than GP destabilization that remain to be identified (35).

**Fig 4:**
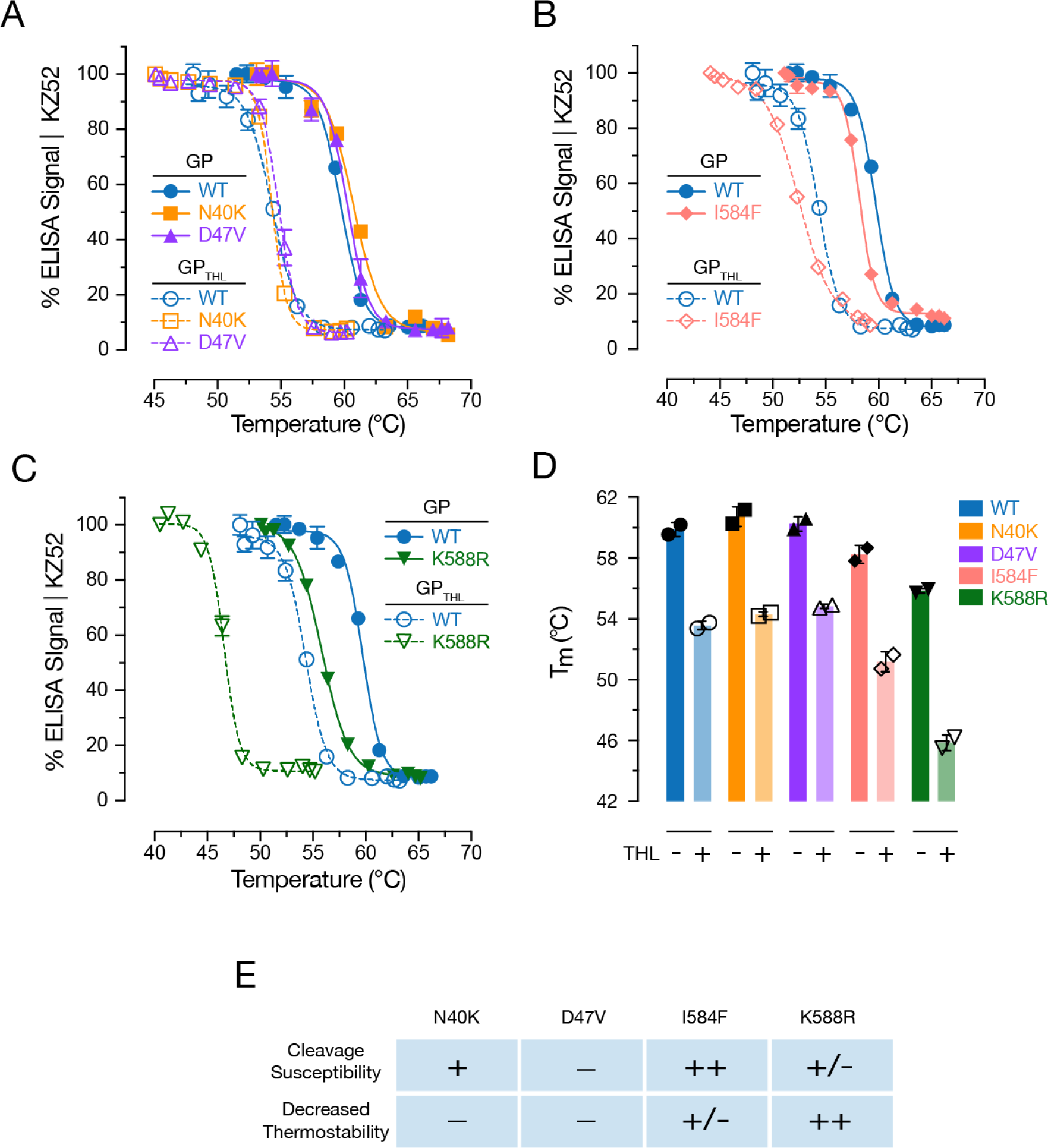
Thermostability of EBOV GP mutants with altered proteolytic requirements for viral entry. Thermal denaturation curves for GP and GP_THL_ for CatB-independent GP mutants N40K and D47V (A), I584F (B) and K588R (C) relative to WT. Averages±SD, n=6 from 2 independent experiments. WT EBOV curves (Blue) from (A) are shown for comparison in (B) and (C). (D) Average T_m_ values from two independent experiments. (E) Summary of the thermostability (A–D) and protease sensitivity phenotypes (from (44)) for the indicated GP mutants.

### Endosomal acid pH stabilizes GP’s pre-fusion conformation

Endosomal acid pH is critical for filovirus entry and appears to play multiple roles in this process: previous work implicates it in the activity of the cysteine cathepsins that proteolytically cleave GP (22, 23, 52, 53), for optimal GP-NPC1 binding (54), to induce rearrangement of the GP2 fusion loop to a membrane-active form (55), and to stabilize the GP2 post-fusion six-helix-bundle structure (52, 56). In addition, acid pH has been proposed to act as a trigger for viral membrane fusion (36, 55). We reasoned that if acid pH was indeed directly involved in GP fusion triggering, it may be expected to reduce the thermostability of GP’s pre-fusion conformation. Unexpectedly, we observed significant enhancements in the thermostability of both GP and GP_CL_ at pH values below 6.5, with T_m_ value increases of ≍4°C at pH 5.5–6.0 relative to those at pH 8.0 (Fig 5). These findings suggest that acid pH is unlikely to play a direct role in triggering GP_CL_ for viral membrane fusion. Instead, they raise the possibility that the acid-dependent stabilization of GP and GP_CL_ is an adaptive response to endo/lysosomal conditions during entry.

**Fig 5:**
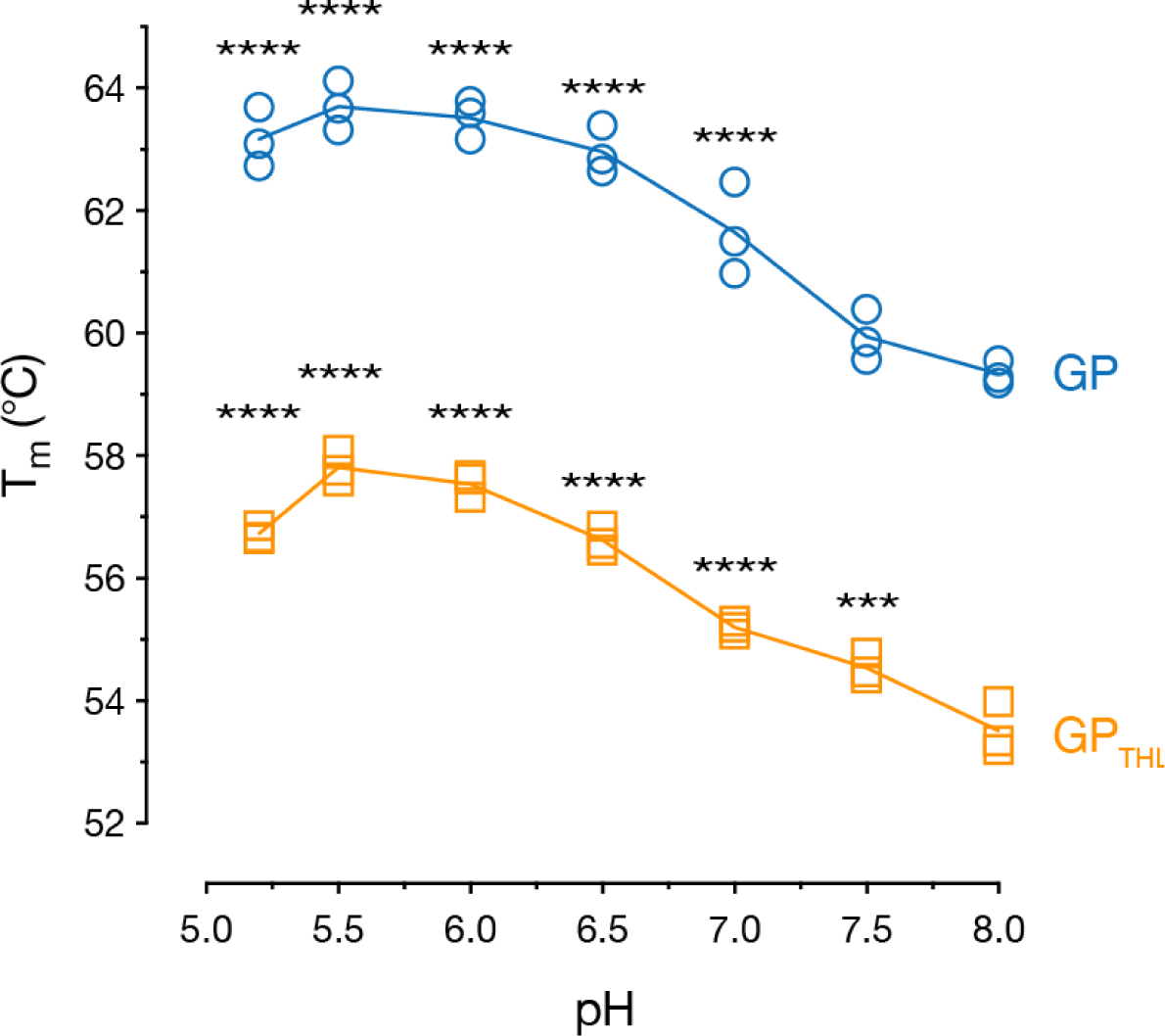
Effect of pH on the thermostability of EBOV GP. rVSVs bearing GP and GP_THL_ were incubated in at the indicated pH values for 1 h at room temp, and then shifted to the indicated temperatures. Virions were then captured onto ELISA plates, and GP was detected at neutral pH using mAb KZ52. The T_m_ values computed from three independent thermal denaturation curves at each pH (n=9) are shown. One-way ANOVA with Dunnet’s post-test was used to analyze the T_m_ relative to pH 8 (***, p<0.001; ****, p>0.0001).

### Small-molecule inhibitor toremifene decreases GP thermostability at acid pH

The SERM toremifene has been proposed to inhibit EBOV infection by destabilizing GP (32). To evaluate the potential destabilizing effect of toremifene in the epitope-loss ELISA, we pre-incubated rVSV-EBOV GP with toremifene and heated viral particles in the presence of the inhibitor at different pH values. As above (Fig 5), the T_m_ was observed to increase with decreasing pH for both GP and GP_CL_, with a maximal increase of 4℃at pH 5.7 (Fig 6A). Although toremifene had no effect on GP thermostability at mildly alkaline pH (7.5–8.0), it decreased GP thermostability relative to the vehicle control at acidic pH values, with a maximal decrease of about 4℃at pH 5.2 (Fig 6A). By contrast, toremifene had a smaller effect on the thermostability of GP_THL_ (Fig 6A). We next tested the dose dependence of toremifene’s capacity to destabilize GP at pH 7.5 and 5.2. Drug concentrations greater than 3.3 µM were necessary for toremifene-mediated GP destabilization, with significantly greater effects seen at acid pH, as described above (maximal ΔT_m_ of ∼9°C at pH 5.2 vs ∼2°C at pH 7.5) (Fig 6B). These findings are in line with those reported by Zhao and co-workers using a recombinant GP ectodomain in a completely different (fluorescence-based) thermostability assay (32). Together, these results confirm that toremifene destabilizes EBOV GP in an acid pH- and dose-dependent manner.

**Fig 6:**
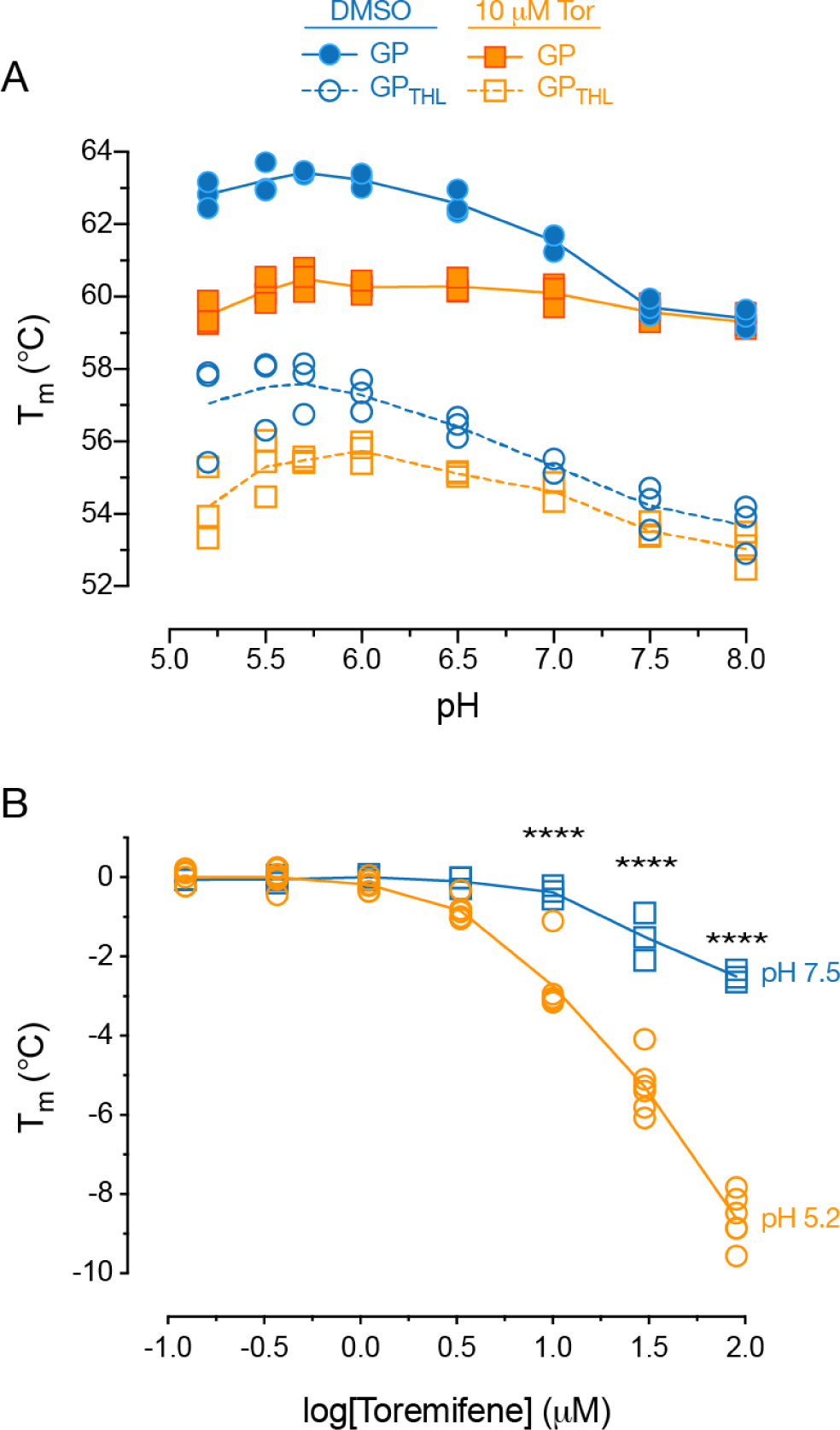
Effect of toremifene on EBOV GP thermostability. (A) rVSV-EBOV GP particles were diluted in PBS adjusted to the indicated pH values containing toremifene (10 µM) or 0.1% DMSO, incubated for 1 h at room temperature, and then incubated at the indicated temperatures. Virions were then captured onto ELISA plates, and GP was detected at neutral pH using mAb KZ52. The T_m_ values computed from three independent thermal denaturation curves at each pH (n=9) are shown. (B) Effect of toremifene concentration on GP thermostability at pH 5.2 vs. pH 7.5 determined as described in panel A. The T_m_ values computed from five independent thermal denaturation curves for pH 5.2 (n=15) and three independent curves for pH 7.5 (n=9) are shown.

### SERM-mediated GP destabilization and inhibition of EBOV infection are not fully correlated

Among the SERMs known to inhibit EBOV entry, only toremifene has been tested for its effect on GP stability (32–34, 57). Here, we tested two additional SERMs and structural analogs of toremifene—clomifene and ospemifene. Entry by VSV-GP_ΔMuc_ was strongly inhibited by toremifene with an IC_50_ of ∼450 nM (Fig 7B), as previously reported (58). Clomifene and ospemifene were much less potent, with IC_50_s of 2.6 µM and 8.7 µM respectively (Fig 7B). Indeed, complete entry inhibition was not observed with ospemifene even at the highest non-cytotoxic doses tested. VSV-G infection, as a control, was not affected by any of the drugs, indicating specific inhibition of EBOV GP-mediated entry. Despite their reduced potency as entry inhibitors, clomifene and ospemifene closely resembled toremifene in their capacity to destabilize GP in the epitope-loss ELISA (Fig 7C). This disconnect between the inhibition of GP-dependent viral entry and the destabilization of GP strongly suggests that mechanisms other than GP destabilization account for the potent antiviral activity of toremifene.

**Fig 7:**
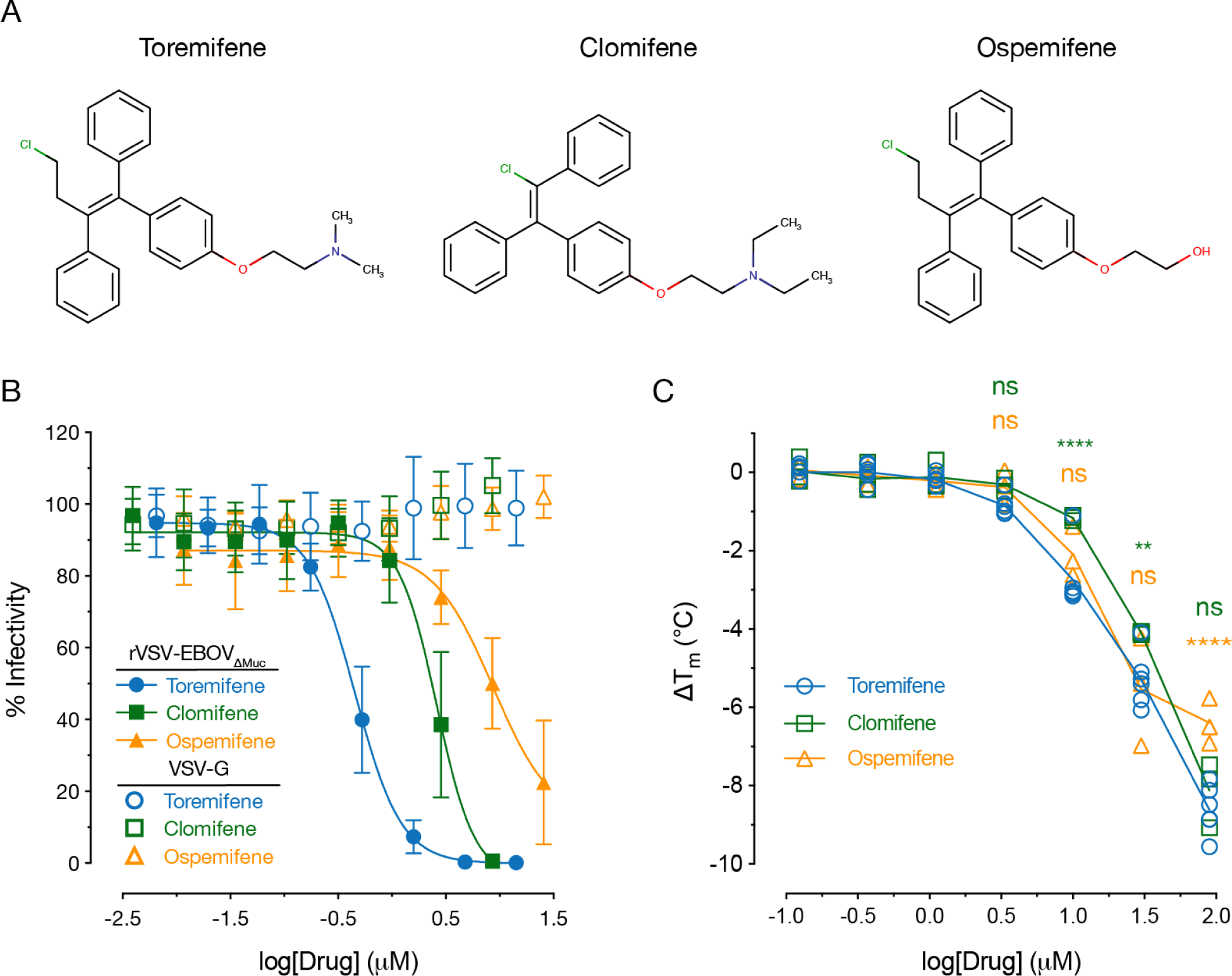
Effect of toremifene structural analogs on EBOV GP-dependent entry and thermostability. (A) Chemical structures for toremifene and two SERMs that are structural analogs, clomifene and ospemifene. (B) SERM-mediated inhibition of rVSV-EBOV GP entry. Averages±SD, n=9 from 3 independent experiments. (C) Effect of clomifene and ospemifene on the thermostability of EBOV GP was determined with KZ52 as described above. Data for the effect of toremifene on thermostability of EBOV GP are from Fig 6 and are shown for comparison. T_m_ values computed from three independent thermal denaturation curves at each pH (n=9) are shown.

### Reduced susceptibility of MARV GP to toremifene-mediated destabilization and viral entry inhibition

Previous work indicates that Marburg virus (MARV entry is inhibited by toremifene, but to a lesser degree than is EBOV (58, 59). We were able to confirm these findings (Fig 8A). Whether toremifene also affects MARV GP stability has not been reported, however. Accordingly, we used MR191, a human mAb specific for the NPC1-binding site in MARV GP (60), to adapt the epitope-loss ELISA to rVSV-MARV GP. MARV GP was significantly less thermostable than EBOV GP (Fig 8B and C). Further, acid pH increased the T_m_ of MARV GP to an extent similar to that of EBOV GP, suggesting that these divergent glycoproteins share at least some molecular determinants of acid-dependent stability (Fig 8A and B). We next examined the thermostability of MARV GP in the presence of toremifene. Concordant with rVSV-MARV GP’s reduced susceptibility to toremifene (Fig. 8A), MARV GP was also less sensitive than EBOV GP to toremifene’s destabilizing effects (Fig 8D). Our findings strongly suggest that toremifene can bind to and destabilize MARV GP, despite substantial differences between MARV and EBOV GPs in the configuration of the toremifene-binding pocket (Fig 9).

**Fig 8:**
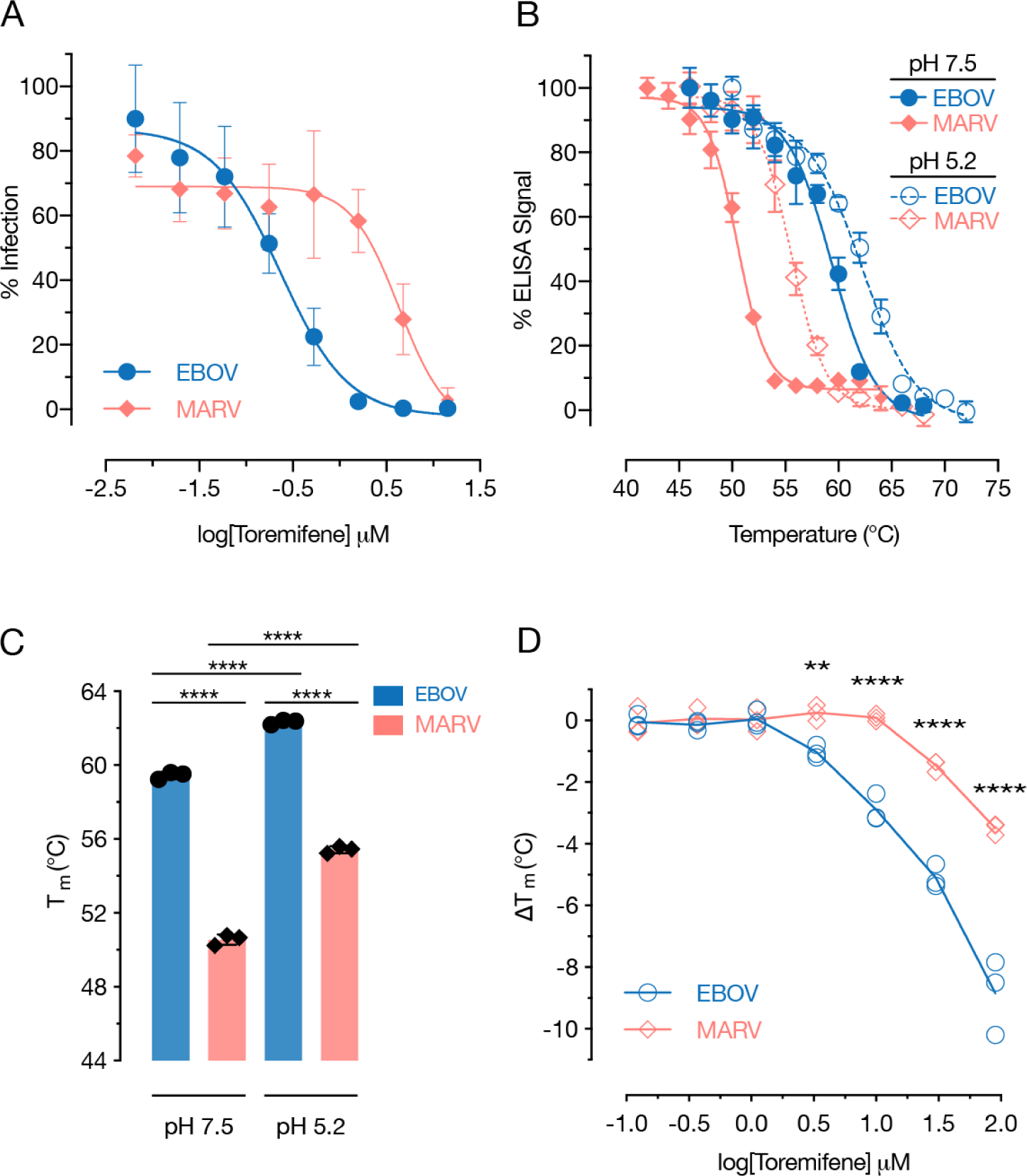
Thermostability of MARV GP and its susceptibility to toremifene. (A) The effect of toremifene on rVSV-MARV GP in Vero cells was determined as described in Fig. 7B. Averages±SD, n=9 from 3 independent experiments. (B–C) Thermal denaturation curves for rVSVs MARV and EBOV GP were determined at pH 7.5 and pH 5.2 with MR191 (MARV) and KZ52 (EBOV), respectively. Averages±SD, n=9 from 3 independent experiments. (C) Calculated T_m_ values from the thermal denaturation curves in panel B. Averages±SD, n=3 from 3 independent experiments. ****, p<0.0001 by One-way ANOVA with Tukey’s post-test correction for multiple comparisons. (D) Effect of toremifene on the thermostability of MARV and EBOV GP at pH 5.2 was determined as described above. The T_m_ values computed from three independent thermal denaturation curves at each pH (n=9) are shown. EBOV vs. MARV: **, p<0.01; ****, p<0.0001 by two-way ANOVA with Sidak’s post correction for multiple comparisons.

**Fig 9:**
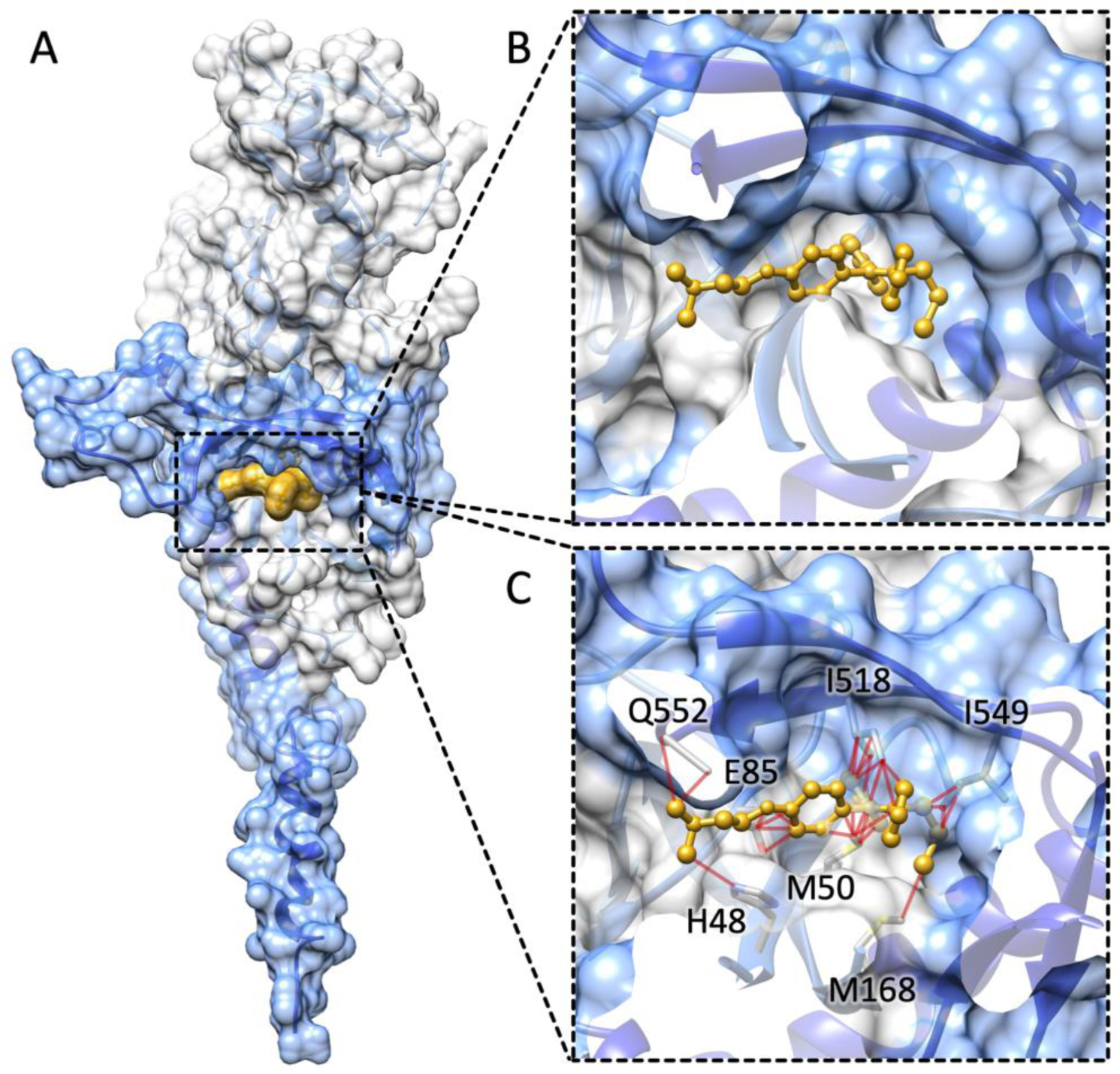
Docking by structural superposition reveals steric clashes between toremifene and MARV GP in the binding pocket. (A) Overview of the EBOV GP monomer bound to toremifene (GP1: light grey surface, GP2: blue surface, toremifene: yellow surface; PDB 5JQ3) (32). (B) Closeup view of toremifene (yellow ball-and-sticks) bound to EBOV GP (GP1: light grey surface, GP2: blue surface), no steric clashes were observed between the protein and the ligand. (C) MARV GP monomer (GP1: light grey surface, GP2: blue surface; PDB 5UQY) (69) was structurally superimposed to the EBOV GP-toremifene complex (only toremifene is shown as yellow ball-and-sticks) using Chimera (70). Structural clashes (red edges) between atoms in MARV GP (depicted as sticks) and toremifene were identified using Chimera and default parameters.

## Discussion

The thermostability of viral glycoproteins can reflect their fusogenicity (27–29). To extend previous studies of filovirus GP thermostability, which rely on purified, recombinant GP ectodomains, we developed and characterized a simple, readily accessible antibody-based assay that measures the stability of full-length GP trimers embedded in viral membranes (Fig 1 and 2). Our findings were both qualitatively and quantitatively concordant with the results of previously published GP thermostability assays (32, 36). One advantage of the approach we have developed is that it allows us to probe the thermostability of specific sequences in GP; by contrast, the fluorescence-based thermal shift assays typically employed for this purpose do not provide such region-specific information. Herein, we found that the highly structured subdomains of the GP trimer—the base and glycan cap—both undergo a concerted loss of conformation at a characteristic temperature range (Fig 1C), whereas the GP stalk, comprising sequences in the GP2 HR1 and HR2 heptad repeat-forming sequences, is relatively refractory to thermal denaturation (Fig 1C). The availability of epitopes in the Muc domain actually increased at elevated temperatures, consistent with previous work suggesting it is intrinsically disordered (Fig 1D).

Another advantage of our approach is its capacity to interrogate the thermostability of divergent GPs through the use of engineered rVSVs and conformation-sensitive mAbs with pan-ebolavirus and pan-filovirus reactivity. Here, we showed that the GPs of the newly discovered ebolavirus BOMV and the marburgvirus MARV are much less stable than those of EBOV and SUDV (Fig 3B-C and 8B-C) suggesting differences in their entry mechanisms that remain to be defined. We speculate that variations in GP thermostability among filoviruses (with attendant consequences for route/efficiency of cell entry and environmental stability) may impact their capacity to infect different types of hosts and spread between them in nature.

Using this thermostability assay, we investigated the effect of GP proteolytic priming—an essential step in filovirus entry—on EBOV GP stability. Our results, together with previous findings (36, 44), indicate that removal of the glycan cap and cleavage within the partially disordered β13–14 loop connecting the base and glycan cap sharply reduce thermostability (Fig 2), whereas the removal of Muc has little effect (Fig 1A). Interestingly, differences in the C–terminus of the cleaved GP1 subunit generated by THL and CatL did not significantly impact GP_CL_ thermostability (Fig 2B-C and 2D-E). Because the additional processing of GP1_CatL_ at its C–terminus by CatB (mimicked by THL) is required for entry (23, 44), we infer that the role of this latter cleavage step is likely not to destabilize GP *per se* but to prime GP for the action of another, unidentified entry host factor (35).

To further investigate the importance of GP destabilization by proteolytic cleavage during entry, we analyzed the thermostability of our previously described CatB-independent GP mutants (Fig 4) (44). Combined with their known protease sensitivity phenotypes (44), our analysis of thermostability suggests three distinct mechanisms of CatB independence (summarized in Fig 4E). Only one of the mutants, GP(K588R), which is slightly more susceptible to proteolysis, exhibited a strong reduction in thermostability that may afford bypass of the requirement for cleavage-mediated destabilization (Fig 4C). More protease-sensitive mutants, including GP(I584F) and to a lesser extent, GP(N40K), possessed WT thermostability (Fig 1A and B). We postulate that these mutations confer CatB independence by accelerating cleavage by other endosomal cysteine cathepsins, such as CatL. The mechanism by which D47V, which alters neither thermostability (Fig 1A) nor protease sensitivity (44), affords CatB independence is presumably distinct from the above. Thus, although thermostability can correlate with CatB independence, increased protease sensitivity and decreased thermostability at least partly reflect distinct molecular mechanisms by which GP proteins can bypass the CatB cleavage requirement during entry.

These apparently complex relationships among the molecular bases of GP thermostability, protease sensitivity, and CatB dependence also extend to other filoviruses. As reported previously, many filovirus GPs, including those of SUDV and MARV, are CatB-independent (24, 47). Although filovirus GPs are indeed polymorphic at some of the amino acid sequence positions altered in the CatB-independent EBOV GP mutants, the mutation of SUDV or RESTV GP to the cognate EBOV GP residues at these positions did not render them CatB-dependent (24), suggesting the existence of other unknown molecular determinants. Similarly, we observed that the CatB independence of SUDV GP could not be explained by its thermostability (Fig 3B and C), since it resembled EBOV GP in this regard. We did find, however, that MARV GP, which is CatB-independent, is much less thermostable than EBOV GP (Fig 8B and C), raising the possibility that both of its phenotypes share a molecular basis similar to that of EBOV GP(K588R). More work is needed to test this hypothesis and to determine if BOMV GP, which is much less stable than EBOV GP (Fig 3B and C), is also CatB-independent.

Given the many roles endosomal acid pH is proposed to play in filovirus entry, we measured the pH dependence of GP thermostability and found, counterintuitively, that acid pH, stabilizes both uncleaved and cleaved GP. This effect appears to be broadly shared among filoviruses, including the divergent EBOV and MARV GPs (Fig 5 and Fig 8B-C). Further, the common behavior of GP and GP_CL_ indicates that the sequences that modulate the pH dependence of GP thermostability reside within its base or stalk subdomains and not the glycan cap or Muc. Our findings complicate previous hypotheses proposing that acid pH serves as part of the molecular trigger for viral membrane fusion (36,55,61): if this were the case, one might expect GP stability to be reduced at acidic pH values, as reported for other acid-triggered Class I viral membrane fusogens (27). We speculate that the acid pH-mediated stabilization of GP plays a functional role in filovirus entry. It may, for instance, help to explain the increased affinity of GP for NPC1 at acid pH (54). Alternatively or in addition, it may prevent premature GP2 fusion loop deployment until viral delivery to late endo/lysosomal compartments, where NPC1 binding and/or additional GP cleavage events can drive fusogenic conformational changes (35). Finally, these observations may also explain the behavior of toremifene and some other EBOV entry inhibitors, which selectively destabilize GP at acid pH (Fig 6 and 7D) (32, 32, 34). We propose that toremifene exerts its antiviral effect, in part, by counteracting the acid-dependent stabilization of GP_CL_ and promoting premature GP_CL_ triggering in endosomal compartments.

Although toremifene can potently block EBOV entry, it only modestly inhibits MARV entry (Fig 8A) (58, 59). Using our epitope-loss ELISA, we found that toremifene is a less potent destabilizer of MARV GP than EBOV GP at acid pH (Fig 8D), concordant with its reduced antiviral activity. A structural alignment of the toremifene-bound EBOV GP and apo MARV GP X-ray crystal structures (Fig 9)) suggests that the geometry of the putative toremifene-binding pocket in MARV GP is not compatible with the EBOV-binding configuration of toremifene, potentially resulting in an alternative binding mode of the drug and a reduction in binding affinity (Fig 9B). Hence. toremifene analogs specifically engineered to fit into the unique pocket in MARV GP may afford enhanced antiviral activity.

Finally, we used viral infectivity assays and the epitope-loss ELISA to more closely examine the relationship between the antiviral activity of toremifene and its analog SERMs, and their capacity to destabilize GP. We found a disconnect between the two phenotypes that is most clearly evident for the close structural analog ospemifene and suggests multiple mechanisms of toremifene action. Specifically, whereas ospemifene resembled toremifene in its capacity to destabilize EBOV GP at acid pH, ospemifene was a much less potent viral entry inhibitor. Given that the two compounds differ only in the substitution of the tertiary amine in toremifene with a hydroxy group in ospemifene, we speculate that they differ not in their capacity to bind to GP, but rather, in the enhanced lysosomotropic activity of toremifene. This property, conferred by toremifene’s tertiary amine, is observed in a large class of otherwise structurally unrelated Class II cationic amphiphilic drugs (CADs) (25, 62–65), and is expected to enhance the accumulation of toremifene, but not ospemifene, in acidic intracellular compartments. Concordantly, we have observed that toremifene induces profound changes in the morphology and dynamics of cellular endo/lysosomal compartments (Mittler and Chandran, manuscript in preparation). These unwanted effects may limit the utility of toremifene and other CADs as anti-filovirus therapeutics.

In sum, we describe a simple and highly adaptable assay that can be used to measure the thermostability of viral membrane-embedded GP proteins from diverse filoviruses under a variety of conditions relevant to cell entry. Aside from its utility in mechanistic studies of filovirus entry, this assay should facilitate the engineering of GP variants with enhanced stability for use in vaccine development, the discovery of new antiviral drugs that alter GP stability, and the identification of host factors that drive or inhibit filovirus entry by modulating GP fusogenicity.

## Methods

### Cell lines and viruses

Vero African grivet kidney cells were cultured in Dulbecco’s modified Eagle medium (DMEM) (Life Technologies, Carlsbad, CA) supplemented with 2% fetal bovine serum (Atlanta Biologicals, Flowery Branch, GA), 1% penicillin streptomycin (Life technologies) and GlutaMAX. Cells were grown at 37℃and 5% CO_2_ in a humidified incubator.

The recombinant vesicular stomatitis Indiana viruses (rVSV) encoding eGFP in the first position with GP proteins from EBOV/Mayinga (EBOV/H.sap-tc/COD/76/Yambuku-Mayinga), SUDV/Boneface (SUDV/C.por-lab/SSD/76/Boneface), and BOMV (BOMV/Mops condylurus/SLE/2016/PREDICT_SLAB000156) in place of VSV G were generated as previously described (44, 51, 66). Viruses containing EBOV/Mayinga lacking Muc, rVSV-GP_ΔMuc_, were generated by genetic deletion of residues 309-489 in the rVSV vector (37). Viruses encoding CatB-independent mutant EBOV GPs were generated previously (44).

To generate rVSV bearing MARV GP, the a previously described vector, VSV(mNG-P)ΔG (66, 67) encoding the fluorescent protein mNeonGreen (mNG) fused to the VSV phosphoprotein (P) was engineered to encode MARV GP (MARV/H.sap-tc/KEN/80/Mt. Elgon-Musoke) in the VSV G position. rVSV(mNG-P)-MARV GP was recovered using a plasmid-based rescue system in 293T cells as described previously (68) and amplified in Vero cells. Viral genomic RNA isolated from viral supernatants was subjected to RT-PCR with VSV genome-specific primers flanking the *GP* gene as previously described, and the sequence of the resulting cDNA was verified by Sanger sequencing.

All experiments with rVSVs were carried out using enhanced biosafety level 2 procedures approved by the Einstein Institutional Biosafety Committee.

### Antibodies

ADI-15878, ADI-15750, and ADI-16061 were described previously (39). KZ52 (38) was kindly provided by Dennis Burton (Scripps Research Institute, La Jolla, CA). MR191 (45) was kindly provided by Zachary Bornholdt (Mapp Biopharmaceuticals, San Diego, CA, USA). MR72 (45) was expressed and purified as described previously (66). 14G7 (40) was kindly provided by John Dye (USAMRIID, Fort Detrick, MD, USA).

### *In vitro* proteolytic cleavage reactions

Cleavage conditions, including enzyme concentrations and cleavage times, were optimized for complete cleavage of GP→GP_CL_ by western blot as indicated below. rVSV-EBOV GP and GP_ΔMuc_ were cleaved with 500 ng/µL thermolysin (THL; Sigma Aldrich, St. Louis, MO) for 60 min and 30 min, respectively. rVSV-SUDV GP and rVSV-BOMV GP were cleaved with THL (250 ng/µL for 60 min). The CatB-independent mutant EBOV GPs were cleaved with THL using previously described and optimized conditions for complete cleavage (200 ng/uL for 60 min) (44). THL cleavage reactions were terminated by adding 10 mM phosphoramidon (Peptides International, Inc., Louisville, KY). For the CatL cleavage experiments, recombinant human CatL (R&D Systems Inc., Minneapolis, MN) was activated on ice for 30 min, then rVSV-EBOV GP was cleaved for 60 min (2 ug/mL) at pH 5.5; reactions were terminated with 0.1 mM E-64 (Sigma Aldrich, St. Louis, MO), a broad-spectrum cysteine protease inhibitor. All THL and CatL reactions were conducted at 37°C. Following cleavage, viruses were immediately used for ELISA experiments as described below.

### SDS-PAGE and western blotting

rVSVs bearing uncleaved, THL-cleaved, or CatL-cleaved GPs were incubated with Protein N– glycosidase F (PNGaseF, 250U; New England Biolabs, Ipswich, MA) under reducing conditions for 16 h at 37℃to remove N–linked glycans. Deglycosylated samples were then resolved in 10% Novex Tricine SDS-polyacrylamide gels (ThermoFisher, Grand Island, NY). rVSV-BOMV GP and THL-cleaved GPs were instead resolved in Bolt 4–12% gradient gels (ThermoFisher), with EBOV GPs for comparison. GP was detected by western blotting with an anti-GP1 polyclonal rabbit serum described previously (22) followed by an Alexa-Fluor 680 dye conjugated anti-rabbit secondary antibody (ThermoFisher). Blots were imaged using the LI-COR Fc fluorescence imager (LI-COR, Lincoln, NE)

### Epitope loss ELISA

rVSVs were cleaved as indicated above. Cleaved and/or uncleaved virions were diluted in PBS (pH 7.5) and then incubated at a temperature range from 42 to 80°C in a thermocycler (actual range for each virus indicated in Results) for 10 min followed by a temperature ramp to 4°C. After cooling, virus was directly captured onto high-binding 96 well half-area ELISA plates (Corning, Corning, NY). Plates were then blocked using 3% BSA in PBS. EBOV GP was detected with KZ52, ADI-15878, ADI-15750, ADI-16061, MR72, or 14G7 as indicated. MARV GP was detected with MR191. Bound antibody was detected with an anti-human antibody conjugated to horseradish peroxidase (HRP; EMD Millipore, Burlington, MA) and Ultra-TMB substrate (ThermoFisher). All binding steps were carried out at 37°C for 1 h. Binding curves were generated using Prism (GraphPad Software, La Jolla, CA) (non-linear regression, variable slope [four parameters]).

The effect of acid pH on GP thermostability was assessed by ELISA as above, except that the PBS incubation buffer was adjusted to pH values ranging from 5.2 to 8.0 as indicated during the heating step. To assess the effect of toremifene on GP thermostability, virus was pretreated with the drug for 1 h at room temperature and heated in the presence of 10 µM toremifene in PBS adjusted to pH ranging from 5.2 to 8.0. Drug dose-response experiments were conducted by pretreating virus with toremifene citrate, clomifene citrate (Sigma Aldrich), ospemifene (Santa Cruz Biotechnology, Dallas, TX) or 1% DMSO as vehicle control for 1 h at pH 5.2 and room temperature. Here, EBOV GP was detected with KZ52 and MARV GP with MR191, respectively. In these experiments, all antibody incubations were carried out at pH 7.4 to eliminate possible pH effects on antibody binding.

### Control ELISAs to verify virion and protein Capture

rVSV-EBOV GP_ΔMuc_ were membrane-labeled by incubating virions with 5 mM function-spacer-lipid (FSL)-biotin (Sigma Aldrich, St. Louis, MO) for 1 h at 37℃as described previously (47). In separate reactions, GP was directly labeled using the amine-specific biotinylation reagent EZ-Link NHS-PEG_4_-Biotin (ThermoFisher), according to the manufacturer’s instructions. Virions were then incubated at a range of temperatures and bound to ELISA plates as indicated above. Biotin-labeled virions were detected using Pierce Streptavidin conjugated to HRP (Strep-HRP; ThermoFisher), followed by detection with Ultra-TMB substrate.

### rVSV infections

VSVs bearing EBOV GP_ΔMuc_ or VSV G were incubated with decreasing concentrations of toremifene, clomifene, or ospemifene in DMEM for 1 h at room temperature. The virus-drug mixture was added to confluent Vero cells in 96 well culture plates, and incubated for 1 h at 37°C. To avoid additional rounds of infection, 20 mM NH_4_Cl was added. Percent infection was scored 14–16 hrs p.i. using Cytation 5 cell imager (Biotek, Winooski, VT). Infection was normalized to control (0 µM drug) and percent infectivity curves were generated using Graphpad Prism (non-linear regression, variable slope [four parameters]).

### Chemical Structures

Small molecule inhibitor structures were prepared using MarvinSketch 19.9.0, 2019, ChemAxon (http://www.chemaxon.com).

## Acknowledgements

We acknowledge I. Gutierrez, E. Valencia, and L. Polanco for laboratory management and technical support. We thank J.M. Fels for his comments and input on earlier versions of the manuscript. This work was supported by National Institutes of Health (NIH) grant R01AI134824 (to K.C.) and the USAID PREDICT program GHN-A-OO-09-00010-00 (to S.J.A.). R.B.III. was partially supported by NIH training grant 2T32GM007288-45 (Medical Scientist Training Program) at Albert Einstein College of Medicine.

